# Quality of Whole Genome Sequencing from Blood versus Saliva Derived DNA in Cardiac Patients

**DOI:** 10.1101/532564

**Authors:** Roderick A. Yao, Oyediran Akinrinade, Marie Chaix, Seema Mital

## Abstract

Whole-genome sequencing (WGS) is becoming an increasingly important tool for detecting genomic variation. Blood derived DNA is the current standard for WGS for research or clinical purposes. We compared the level of microbial contamination, sequencing coverage, as well as yield and concordance of single-nucleotide polymorphism (SNP) and copy number variant (CNV) calls in WGS from paired blood and saliva samples from 5 pediatric heart disease patients. We found that although saliva samples contained a higher proportion of sequence reads that map to the human oral microbiome, these reads were readily excluded by mapping the reads to the human reference genome. Sequencing coverage was low only in 1 of 5 saliva samples. Over 95% SNPs (including rare SNPs) but <80% CNVs called in blood genomes were detected in paired saliva genomes. These findings suggest that most good quality saliva samples can serve as an alternative to blood samples for detection of sequence variants from WGS in cardiovascular disease patients.

## INTRODUCTION

As the cost of sequencing continues to decrease, whole genome sequencing (WGS) is being increasingly used for both research and clinical applications to study the role of genetic variation in human disease. The use of blood derived DNA is the current standard for WGS. Published studies report that saliva-derived DNA can be used for array genotyping (Bahlo et al. 2010) and whole-exome sequencing (Kidd et al. 2014) as long as the quantity of human DNA in each sample is sufficient. Wall et al. (Wall et al. 2014) reported finding no differences in sequencing quality or variant call error rate between blood and saliva samples for both whole exome sequencing (WES) and WGS. However, there have been very few studies comparing WGS results from paired blood and saliva-derived DNA.

To determine whether saliva-derived DNA can serve as an adequate substitute for blood-derived DNA, we compared WGS data from blood- and saliva-derived DNA from pediatric patients recruited into the Heart Centre Biobank Registry, a biorepository for childhood onset heart disease (Fung et al. 2013; Papaz et al. 2012). Specifically, we compared the proportion of sequencing reads that map to non-human sources in blood versus saliva, the sequencing coverage between blood and saliva samples, and the concordance of single nucleotide variant (SNV) and copy number variant (CNV) calls between blood and saliva samples.

## METHODS

### Study Samples

Study participants were derived from the Heart Centre Biobank Registry, a multi-center biorepository that has been prospectively enrolling pediatric and adult patients with (or at risk for) heart disease from six institutions across the province of Ontario, Canada since 2007 (Fung et al. 2013). We accessed paired blood and saliva samples from five unrelated individuals participating in the Biobank. Two (sample pairs 1 and 2) were probands diagnosed with tetralogy of Fallot (TOF), a type of congenital heart disease, and three (sample pairs 3, 4 and 5) were probands diagnosed with hypertrophic cardiomyopathy (HCM). All participants and/or their parents or legal guardians provided written, informed consent and the study was approved by the Institutional Research Ethics Boards at each participating site.

### DNA Quality

2-5 ml blood was collected in EDTA tubes and 2-4 ml saliva was collected using Oragene saliva kits. DNA was extracted from blood or saliva through Chemagic Star robotic system using a magnetic bead methodology at the SickKids Centre for Applied Genomics. Quality control checks were performed using Agarose Gel Electrophoresis and Nanodrop 2000 Spectrophotometer to verify DNA integrity. DNA quantification was measured using Qubit 3.0 Fluorometer to confirm DNA concentration. DNA samples were deemed to have met QC thresholds if they had a single clear band on the agarose gel, a minimum DNA concentration of 20 ng/μl, and a 260/280 absorbance ratio greater than 1.3. A total of 1 mcg of DNA (final volume 30 μl) at a minimum concentration of 20 ng/μl was used for WGS.

### Sequencing, Read Alignment, and Variant Calling

WGS was performed using Illumina HiSeq X to a target average coverage depth of 30x. Sequencing read alignment was done using Isaac Aligner to human genome build hg19. Single nucleotide variant (SNV), i.e. single-nucleotide polymorphism (SNP), and small insertion-deletion (indel) calling was performed using Isaac Variant Caller with default parameters. To interpret variant pathogenicity, we implemented an automated variant prioritization pipeline based on the 2015 American College of Medical Genetics and Genomics (ACMG) variant interpretation criteria (Richards et al. 2015). SNVs identified as pathogenic or likely pathogenic by the pipeline were manually confirmed for pathogenicity. Copy number variants (CNVs) were called using Control-FREEC (Boeva et al. 2012) for Sample Pairs 1 and 2 or Canvas (Roller et al. 2016) for Sample Pairs 3, 4, and 5.

### Downsampling

In order to account for possible bias resulting from differing numbers of reads between blood and saliva samples, each sample was randomly reduced to 730 million reads using samtools (version 1.4.1) (Li et al. 2009). All subsequent analyses on each pair of blood and saliva sequencing datasets were performed in complete datasets and in downsampled datasets.

### Gene Sets and Regions

We compiled a list of 854 cardiovascular disease-associated (CVD) genes which included genes represented in commercially available cardiovascular disease gene panels and previously published genes known to be associated with congenital heart disease, cardiomyopathy, and other cardiovascular diseases (Andersen et al. 2014). The genomic regions covered by the canonical transcripts for CVD genes, as well as for all genes, were obtained from the Consensus CDS (CCDS) database (Farrell et al. 2014; Harte et al. 2012; Pruitt et al. 2009) (see **Supplemental Table S1** for genomic positions of all CCDS transcripts and **Supplemental Table S2** for genomic positions of CVD gene transcripts). For any CVD genes lacking transcripts in the CCDS database, we used the transcript start and end positions from Ensembl GRCh37 release 93 (Zerbino et al. 2018).

### Statistical Analysis

#### Microbial contamination analysis

To find the extent of microbial contamination, we extracted the set of reads from each sample marked as unmapped to the hg19 reference genome using samtools (version 1.4.1) (Li et al. 2009). Then, using FastQ Screen (version 0.11.4) (Wingett and Andrews 2018) running BWA (version 0.7.15) (Li and Durbin 2009), we re-mapped the unmapped reads to the human reference genome hg19 and the microbial sequences from the Human Oral Microbiome Database (HOMD) (Chen et al. 2010) and compared the proportion of previously unmapped reads that remapped to the human genome versus the microbiome.

#### Coverage comparison

We analyzed genome-wide sequencing coverage for each WGS dataset using the genomecov command from the bedtools toolset (version 2.25.0) (Quinlan and Hall 2010) on the aligned reads. We also used the coverage command from the bedtools toolset to find sequencing coverage within the regions covered by all canonical CCDS transcripts and the canonical CCDS transcripts for the 854 CVD genes. All three analyses provided a coverage profile with the number of nucleotides covered at all sequencing depths for comparison between paired samples. We used the cumulative sum for each coverage profile to determine the number of positions covered at or greater than target depth and generated the curves for the cumulative coverage data using the statistical software R (version 3.5.1) (R Core Team 2018). We calculated the proportion of the genome covered at a minimum 20x coverage. Then, we used a paired t-test on the percentage of the whole genome, CCDS transcripts, and CVD gene transcripts covered at 20x or greater in order to find whether coverage in blood was significantly different from coverage in saliva.

#### Variant comparison

We used the isec command from bcftools (version 1.4.1) (Li et al. 2017) to find the intersections of all SNPs called in each blood and saliva sample pair. We annotated SNPs unique to either blood or saliva with snpEff (version 4.3) (Cingolani et al. 2012) to computationally predict variant effects. We repeated this for SNPs falling within canonical CCDS transcripts and within the canonical CCDS transcripts for the 854 CVD genes. Finally, we repeated each of the previous comparisons for rare SNPs, i.e. those were absent or occurred at a minor allele frequency (MAF) of less than 1% in the Genome Aggregation Database (gnomAD) (Lek et al. 2016). All SNVs, CNVs, and rare SNVs were compared for concordance between paired blood and saliva samples. Where clinical genetic test results had identified a pathogenic variant, we compared the detection rate of these known variants in blood versus saliva samples. In addition, we compared the proportion of variants that were concordant between paired blood and saliva and the types of variants, i.e. variants in exonic, intronic, intergenic, pseudogene regions, or causing gene fusions between blood and saliva. In order to find the concordance between CNVs called in each blood and saliva sample pair, we used the intersect command from bedtools (version 2.25.0) (Quinlan and Hall 2010) in order to find which CNVs in each sample had >50% overlap with at least one other CNV called in its counterpart.

## RESULTS

Since the launch of the Heart Centre Biobank Registry in 2007, 7,408 participants were recruited. 531 blood and 502 saliva samples from recruited participants had DNA quality assessed prior to WGS for different research projects. Average DNA quality metrics for the blood and saliva samples are shown in Table 1. Overall, DNA from 46% saliva samples failed QC and were not sequenced compared to only 6% QC failure for DNA from blood samples. The DNA quality metrics for the 5 paired blood and saliva samples showed comparable DNA quality between blood and saliva.

**Table 1.** DNA quality metrics for blood and saliva derived DNA

|  | Pooled samples |  | Paired samples |  |
| --- | --- | --- | --- | --- |
|  | Blood DNA | Saliva DNA | Blood DNA | Saliva DNA |
| Number of Samples | 531 | 502 | 5 | 5 |
| Fluorometric DNA Concentration (ng/μL) | 180 | 153 | 314 | 214 |
| 260/280 Absorbance Ratio | 1.84 | 1.8 | 1.84 | 1.74 |
| 260/230 Absorbance Ratio | 1.96 | 1.32 | 1.36 | 1.28 |
| Samples Failing Mandatory Criteria | 32 (6%) | 231 (46%) | 0 (0%) | 0 (0%) |
| Samples Failing DNA Concentration cut-off (< 20 ng/μL) | 0 | 0 | 0 | 0 |
| Samples Failing 260/280 Absorbance Ratio ($\leq 1.3$) | 0 | 0 | 0 | 0 |
| Samples Failing Agarose Gel | 32 | 231 | 0 | 0 |

### Microbial contamination analysis

An average of 95.5% of all WGS reads from blood samples and 82.6% of reads from saliva samples initially mapped to the hg19 human reference genome. Of the unmapped reads, 2.6% from blood and 2.5% from saliva samples re-mapped to the hg19 human reference using FastQ Screen and BWA, 0.09% and 10.7% from blood and saliva respectively mapped to the human oral microbiome, and 1.05% and 1.03% of reads from blood and saliva respectively mapped to both hg19 and the human oral microbiome. Therefore, read mapping to the hg19 human reference genome was effective in excluding most of the reads from the human oral microbiome. The proportion of final mapped and unmapped reads in each sample are summarized in Figure 1. Although a higher proportion of reads from saliva samples mapped exclusively to the human oral microbiome compared to blood samples, this difference was not statistically significant (p = 0.13). Of note, saliva sample 5 had the highest proportion of reads mapping to the human oral microbiome compared to the other samples. After excluding it, the final proportion of reads mapping to hg19 increased to 98.2% in blood and 93.8% in saliva.

**Figure 1.**
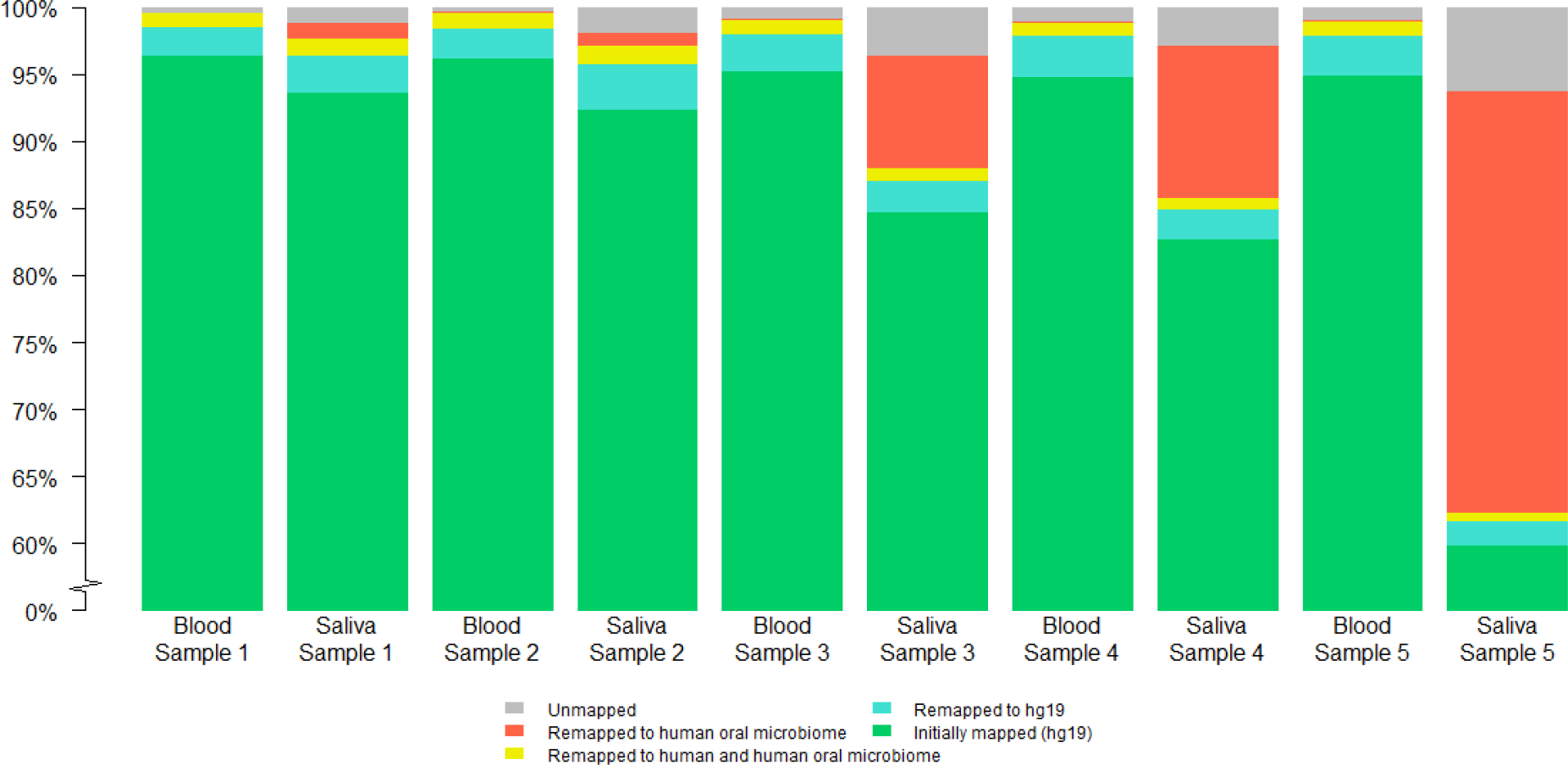
Proportion of reads mapping to each reference genome for each blood and saliva pair

### Coverage analysis

The reported mappable mean depth of coverage was more than 30x for 4 of the 5 paired blood and saliva samples (Table 2). Sample 5 had a lower mean depth of 22.6x. This was seen despite the DNA quality metrics being acceptable in sample 5, which had a DNA concentration of 280 ng/μL, and a 260/280 absorbance ratio of 1.88. Excluding pair 5, the proportion of the genome with at least 20x coverage for all reads ranged from 93% - 96% for blood genomes and from 85% - 94% for saliva genomes (p=0.07). The proportion of the genome with at least 20x coverage for 730M randomly downsampled reads ranged from 91% - 94% for blood genomes and from 84% - 94% for saliva genomes (p=0.19). The proportion of CCDS transcript regions with mean coverage of at least 20x was 94.7% for blood and 92.5% for saliva (p=0.12). The proportion of samples with mean coverage of at least 20x within the subset of transcripts for the 854 CVD genes was 96.9% for blood and 94.6% for saliva (p=0.08). Therefore, saliva samples had overall adequate depth of coverage which was not significantly different from coverage in blood. Figure 2 displays the cumulative coverage for the 5 paired samples across the whole genome, in CCDS transcripts, and in CVD gene regions.

**Table 2.** Sequencing coverage in 5 sample pairs

|  | Sample Pair 1 |  | Sample Pair 2 |  | Sample Pair 3 |  | Sample Pair 4 |  | Sample Pair 5 |  |
| --- | --- | --- | --- | --- | --- | --- | --- | --- | --- | --- |
|  | Blood | Saliva | Blood | Saliva | Blood | Saliva | Blood | Saliva | Blood | Saliva |
| Mappable mean depth | 36.5x | 33.2x | 34.2x | 33.0x | 37.5x | 34.3x | 37.0x | 31.5x | 36.8x | 22.6x |
| Proportion of whole genome with at least 20x coverage for all reads | 96.0% | 94.6% | 95.3% | 94.3% | 93.7% | 85.6% | 93.7% | 89.3% | 93.6% | 63.8% |
| Proportion of whole genome with at least 20x coverage for 730M downsampled reads | 94.5% | 94.5% | 94.3% | 94.3% | 90.8% | 85.6% | 91.1% | 83.9% | 91.0% | 49.8% |

**Figure 2.**
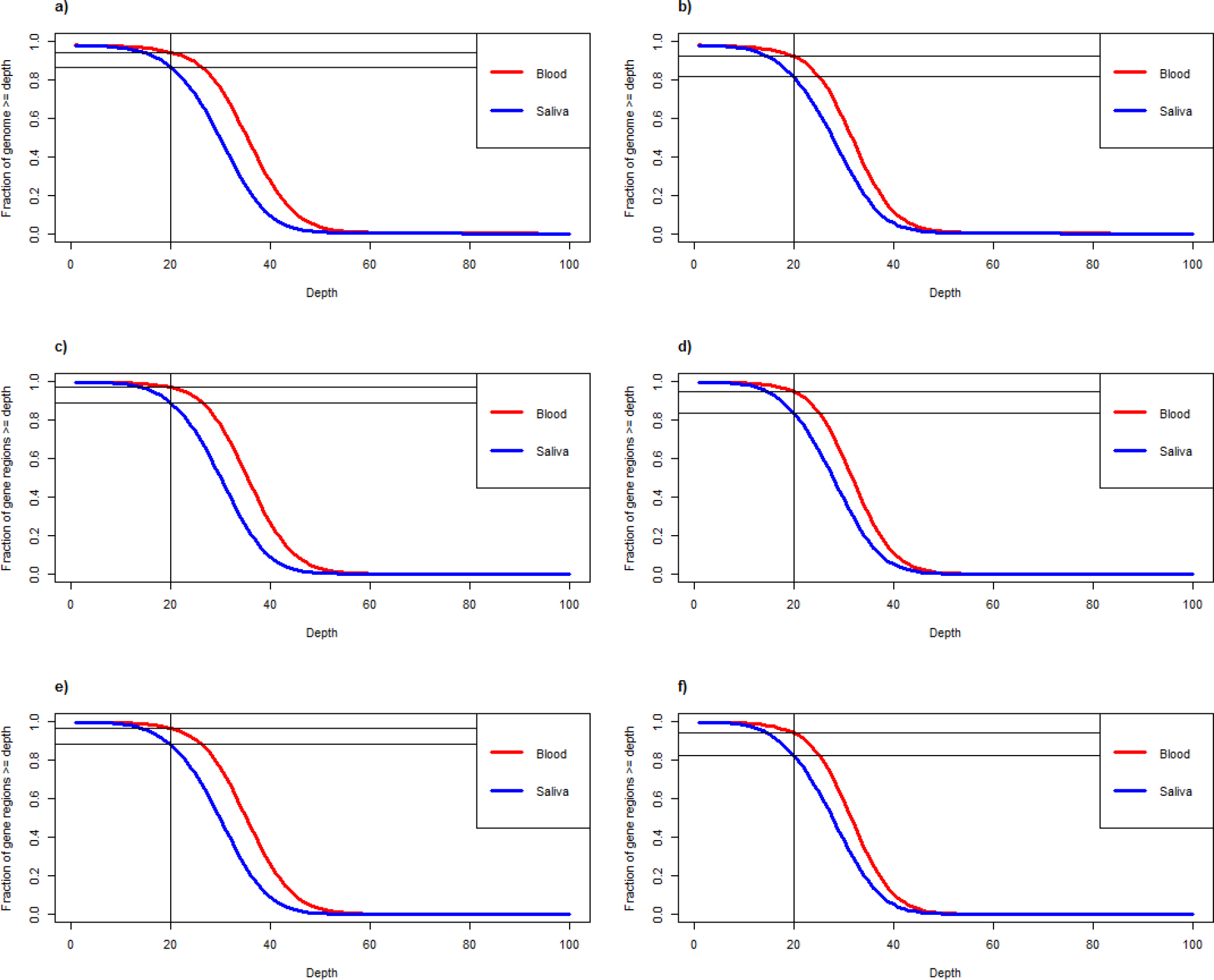
Average sequencing depth curves, displaying the cumulative proportion of the genome covered at a minimum of 20x sequencing depth in 5 paired blood and saliva samples. The vertical lines indicate 20x sequencing depth, while the horizontal lines indicate the fraction of the genome at 20x sequencing depth or greater. (**a**) and (**b**) show the averaged genome-wide coverage curves for all reads and downsampled reads, respectively. (**c**) and (**d**) show the averaged coverage curves for CCDS transcripts for all reads and downsampled reads, respectively. (**e**) and (**f**) show the averaged coverage curves for CVD gene transcripts for all reads and downsampled reads, respectively.

### Variant yield between blood and saliva

#### SNP concordance

In light of a lower proportion of reads from saliva mapping to the human reference genome, we compared if variant yield was also lower in saliva than in blood. The average SNP yield in blood was 3.71M and in saliva was only 1% lower at 3.68M (**Supplemental Table S3**). Also, the proportion of SNPs called in blood that were also detected in the paired saliva sample was >95% in all saliva samples for genome-wide calls, for SNPs in all CCDS transcripts, and for SNPs in CVD genes (Table 3, Figure 3). Of the discordant SNPs, i.e. SNPs unique to either blood or saliva, only 0.6% were exonic; the remainder included 27% intronic, 65% intergenic, and 6.5% within a pseudogene (**Supplemental Table S3)**. There was no difference in the types of discordant SNPs between blood and saliva samples.

**Table 3.** Variant yield in saliva relative to blood genomes

|  | **Sample**<br>**Pair 1** | **Sample**<br>**Pair 2** | **Sample**<br>**Pair 3** | **Sample**<br>**Pair 4** | **Sample**<br>**Pair 5** | **Average** | **SEM\*** |
| --- | --- | --- | --- | --- | --- | --- | --- |
| All SNPs |  |  |  |  |  |  |  |
| Genome-wide SNPs | 96.80% | 96.3% | 98.4% | 98.2% | 96.6% | 97.3% | 0.4% |
| SNPs in CCDS transcripts | 97.40% | 97.0% | 98.8% | 98.6% | 97.2% | 97.8% | 0.4% |
| SNPs in CVD gene transcripts | 97.40% | 97.0% | 98.8% | 98.6% | 97.1% | 97.8% | 0.4% |
| Rare SNPs |  |  |  |  |  |  |  |
| Genome-wide SNPs | 89.90% | 86.3% | 90.1% | 93.0% | 92.5% | 90.4% | 1.2% |
| SNPs in CCDS transcripts | 93.20% | 90.4% | 93.0% | 94.9% | 94.3% | 93.2% | 0.8% |
| SNPs in CVD gene transcripts | 93.00% | 89.2% | 92.9% | 95.1% | 95.0% | 93.0% | 1.1% |
| CNVs |  |  |  |  |  |  |  |
| Genome-wide CNVs | 88.70% | 86.1% | 77.5% | 71.6% | 58.9% | 76.6% | 5.4% |
| CNVs in CCDS transcripts | 95.30% | 91.6% | 70.6% | 78.4% | 52.1% | 77.6% | 7.8% |
| CNVs in CVD gene transcripts | 95.80% | 83.3% | 86.7% | 68.4% | 57.1% | 78.3% | 6.9% |

\*SEM, standard error of mean

**Figure 3.**
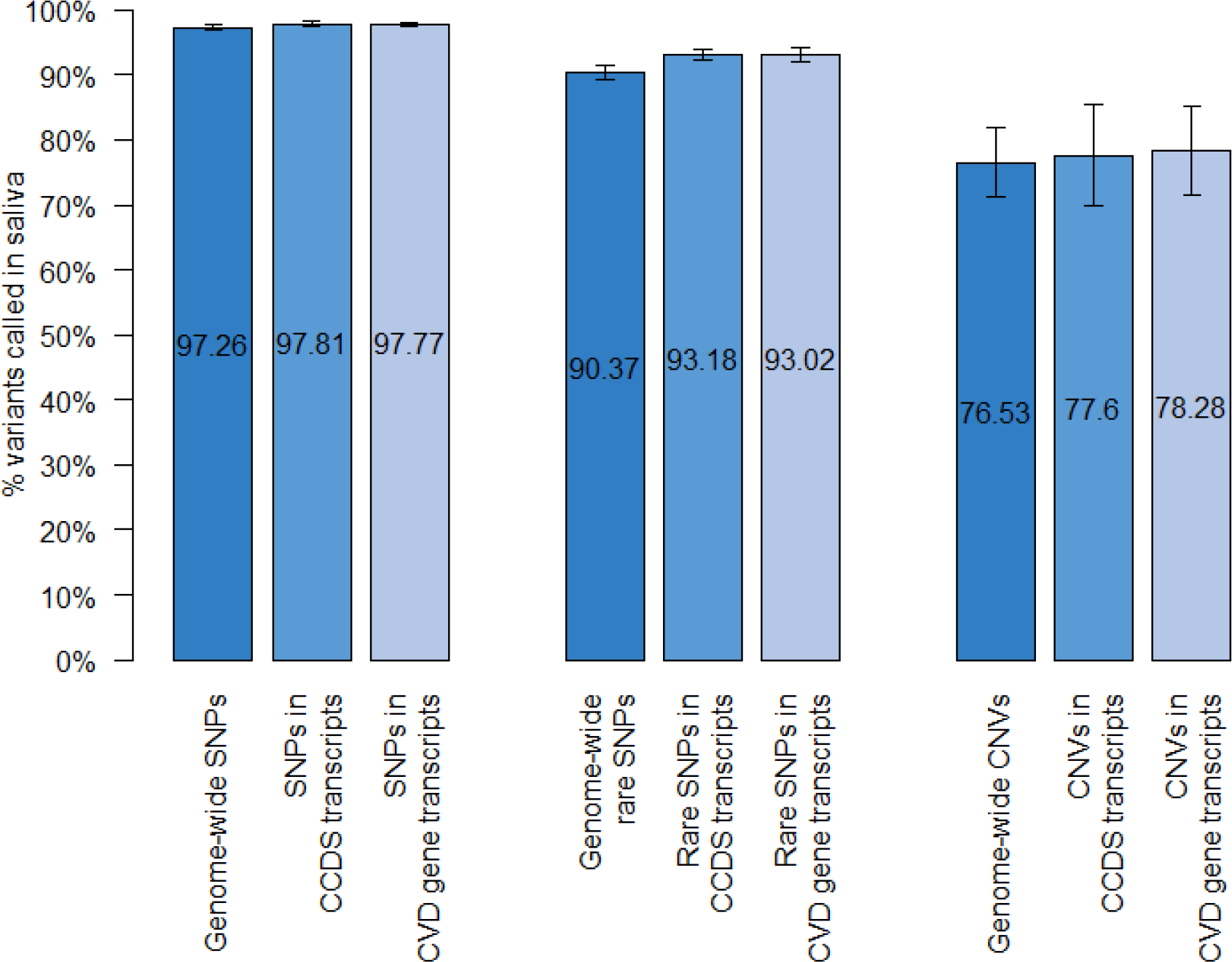
Variant detection in saliva genomes relative to blood genomes (n = 5). The bar graph shows average proportion of variants (SNPs, CNVs) called in blood genomes that were also detected in a paired saliva sample. Rare SNPs were defined as SNPs with a minor allele frequency < 1% in the general population.

Proportion of rare SNPs in blood (MAF<1%) that were also detected in saliva genomes was lower (but over 90% in all samples) at 90.4±3% genome-wide, 93.2±2% in all CCDS transcripts, and 93±2% in CVD genes. Further, rare pathogenic variants and variant/s of uncertain significance (VUS) identified on previous clinical genetic testing in the 3 hypertrophic cardiomyopathy samples were detected in both blood and saliva whole genome pairs from these 3 patients. These included a heterozygous pathogenic missense variant in *MYH7* (NM_000257.4:c.G1208A:p.R403Q) in Sample Pair 3, a heterozygous missense VUS in *MYBPC3* (NM_000256.3:c.T3548G:p.F1183C) in Sample Pair 4, and a heterozygous pathogenic missense variant in *MYH7* (NM_000257.4:c.C2722G:p.L908V) in Sample Pair 5. The comparisons for all SNPs over the entire genome, in CCDS transcripts, and CVD gene transcripts as well as rare SNPs in the aforementioned regions, are described in **Supplemental Tables S4-S9**.

#### CNV concordance

Overall there was considerable variability in CNV calling rates across both blood and saliva samples. The average number of CNV calls were 14% lower in saliva compared to blood. The proportion of CNVs in blood that were also detected in saliva genomes was lower at 76% genome-wide, 77% in all CCDS transcripts, and 78% in CVD genes (Figure 3). The comparisons for all CNVs in each sample across the entire genome, in CCDS transcripts, and CVD gene transcripts including shared and unique CNVs are described in **Supplemental Table S10**. CNV detection was lowest in Sample 5 (<60%) compared to the other 4 sample pairs.

## DISCUSSION

The advances in genomics research has increased the need for collaboration and sample sharing amongst researchers. Population biorepositories often acquire samples from participants remotely. Saliva can be stored for long periods of time and shipped at room temperature which makes sample preservation during shipping easier compared to blood. However, there has been little evidence regarding the suitability of saliva samples for WGS. The study by Wall et al. (2014) found “no difference in the sequencing quality or error rate of blood and saliva samples”, but they did not perform a systematic comparison of the coverage, microbial contamination, or variant concordance in paired blood and saliva samples. Our results showed that a higher proportion of saliva samples (almost 50%) may not meet the stringent QC criteria required for WGS. However for samples that do meet stringent QC criteria, the quality of WGS data from saliva samples is comparable to blood samples in majority of the samples (80% in our study). Our study, using a direct comparison of WGS quality in DNA derived from paired blood and saliva samples, revealed good coverage, low microbial contamination and a high degree of concordance for variant calls, especially sequence variants between blood and saliva samples.

When comparing for specific differences between paired samples, we found higher microbial contamination in saliva-derived DNA, as shown by a higher proportion of short-read mapping to the sequences from HOMD, although this was not statistically different. However, since these reads typically do not map to the human reference genome and remain unmapped during read alignment, they are not included in downstream analyses and therefore are unlikely to have a major effect on variant calling. There remains a concern that reads mapping to the microbiome in saliva samples compete with the mapping of reads to the human genome thereby reducing the resolution of human relevant sequence data. This can be addressed by increasing the target coverage depth with higher resolution sequencing. Our findings suggest that this may not be routinely required for all saliva samples since the WGS quality was comparable to blood in the majority of saliva samples. In particular, rare clinically relevant variants identified on clinical genetic testing were detectable on WGS of saliva and blood samples from the same patients.

We further found that in 4 of the 5 pairs (80%), the proportion of the genome covered at 20x or greater was similar between blood and saliva genomes, with only 2% higher coverage in blood versus saliva at 20x both across the genome as well as within all CCDS transcripts and CCDS transcripts for 854 CVD genes. Of note, Saliva Sample 5 had a higher proportion of reads that mapped exclusively to the human oral microbiome which caused the average read coverage to drop to 22.6x, which is below the average target depth of 30x. The DNA quality from saliva sample 5 was comparable to blood based on agarose gel results, DNA concentration, and 260/280 absorbance ratios. But it is possible that sample 5 was improperly collected with more oral microbiome contamination at the time of collection. In a similar series of comparisons involving whole exome sequencing, Zhu et al. (2015) found that sequences from blood had a 3.3% higher proportion with minimum 20x coverage in blood compared to saliva but this was not significantly different. With randomly downsampling each genomic sequence in order to ensure an equal number of reads between paired samples, there remained a non-significant difference in coverage between blood and saliva genomes, and this difference was lower than the difference reported in exome comparisons by Zhu et al. (2015).

Reassuringly, despite the differences in coverage and microbial contamination in one of the 5 saliva samples, WGS from saliva samples was able to detect 95% of all SNPs detected in a paired blood sample for SNPs genome-wide, within CCDS transcripts, and within CVD gene transcripts. When comparing rare SNPs (MAF < 1%), the proportion of common variants seen in saliva dropped to 90% for all rare SNPs, and 93% for SNPs in CCDS transcripts and in CVD genes with a higher number of discordant SNPs unique to blood or unique to saliva. This limitation needs to be kept in mind when using saliva samples for WGS. Nevertheless, it was reassuring that rare causal SNPs and VUS identified on clinical testing could still be detected by WGS in both blood and saliva samples. There were no systematic difference between functional categories of SNP calls between blood and saliva genomes. CNV yield was more variable across blood and saliva samples with lower proportion of shared CNVs between blood and saliva – 76% for all CNVs, 77% for CNVs in CCDS transcripts, and 78% for CNVs in CVD genes. The lower CNV yield in saliva samples may be the result of the reduced coverage in the saliva sample since both Control-FREEC and Canvas utilize read depth in order to call CNVs (Boeva et al. 2012; Roller et al. 2016). As a result, a reduction in read depth may have a larger effect on CNV calling in a given dataset. On the other hand, the lack of a best practices pipeline for CNV calling (Trost et al. 2018) may also be a contributing factor to the observed inconsistency in CNV calls.

In summary, saliva DNA that met stringent QC checks provided good quality WGS data with comparable detection of common and rare SNPs despite evidence of oral microbiome sequence in some saliva samples but CNV yield in saliva was lower. The utility of saliva samples can be improved by ensuring proper technique for saliva collection, selecting saliva DNA that meets stringent QC criteria, and verifying that target sequencing depth has been met and that at least 90% of reads map to the human genome in WGS data from saliva samples as a minimum threshold for use in downstream analysis. Use of microbiome kit and/or higher depth of sequencing may help WGS yield in samples not meeting QC metrics; however this was not explored in our study. Our findings therefore suggest that good quality DNA from saliva samples can serve as an adequate alternative for WGS when blood samples are not available.

## Data Access

All patients have consented to data sharing, and WGS data files are in the process of being submitted to the European Genome-Phenome Archive (EGA).

## Acknowledgements

We acknowledge the Labatt Family Heart Centre for access to biobanked samples, The Centre for Applied Genomics at the Hospital for Sick Children for providing sequencing services, and Anastasia Miron for providing sample QC data. This work was funded by the Ted Rogers Centre for Heart Research (SM), and the Heart and Stroke Foundation of Ontario Chair in Cardiovascular Science (SM).

